# Rock traits drive complex microbial communities at the edge of life

**DOI:** 10.1101/2021.08.04.455018

**Authors:** Claudia Coleine, Manuel Delgado-Baquerizo, Andrea Zerboni, Benedetta Turchetti, Pietro Buzzini, Laura Selbmann

**Affiliations:** Department of Ecological and Biological Sciences, University of Tuscia, Viterbo, Italy; Departamento de Sistemas Físicos, Químicos y Naturales, Universidad Pablo de Olavide, 41013 Sevilla, Spain; Dipartimento di Scienze della Terra “A. Desio”, Università degli Studi di Milano, Milano, Italy; Department of Agricultural, Food and Environmental Sciences, University of Perugia, Perugia, Italy; Italian Antarctic National Museum (MNA), Mycological Section, Genoa, Italy

**Author notes:** **Corresponding author**: Claudia Coleine; Laura Selbmann.

## Abstract

Antarctic deserts are among the driest and coldest ecosystems of the planet; there, some microbes hang on to life under these extreme conditions inside porous rocks, forming the so-called endolithic communities. Yet, the contribution of distinct rock traits to support complex microbial assemblies remains poorly determined. Here, we combined an extensive Antarctic rock survey with rock microbiome sequencing and ecological networks, and found that contrasting combinations of microclimatic and rock traits such as thermal inertia, porosity, iron concentration and quartz cement can help explain the multiple complex and independent microbial assemblies found in Antarctic rocks. Our work highlights the pivotal role of rocky substrate heterogeneity in sustaining contrasting groups of microorganisms, which is essential to understand life at the edge on Earth, and for searching life on other rocky planets such as Mars.

## Main text

Antarctica supports some of the most unexplored and isolated ecosystems of the planet, wherein existence of life under the most extreme conditions is still being questioned^1^. In the driest and coldest ice-free areas of continental Antarctica (e.g. in the McMurdo Dry Valleys or mountain peaks cropping out from the Polar Plateau), where environmental conditions reach the limits for supporting life endolithic microbes, living within rock surface, often represent the most significant standing biomass, forming self-sustaining ecosystems among which those dominated by lichens are the most organized, widespread and studied^2–4^. We have sound scientific knowledge on the overall role and composition of these rock-colonizing microbes. These pioneer rock microbial communities support key processes such as nutrient cycling, rock weathering, and proto-soil formation, and are composed of simple communities wherein primary producers are lichen-associated or free living chlorophycean algae and cyanobacteria, while consumers are fungi (free-living or lichen symbionts) and bacteria. These studies pushed considerably the general knowledge about their biodiversity, evolution of new and peculiar taxa, and their extraordinary resistance to multiple stresses^5–7^. However, the more specific rock environmental conditions promoting microbial life in these challenging environments, and the capacity of different rock traits to support complex microbial assemblies, are still largely unknown. In the endolithic habitat, micro-environmental conditions inside the rocks may differ significantly from macro-environmental regimes because of the complex interactions between climate and extrinsic and intrinsic rock physico-chemical properties, providing microorganisms with thermal buffering, key micronutrients, physical stability, protection against incident ultraviolet (UV) and solar radiations and, most importantly, ensuring water retention. Herein, we combined rock microbiome sequencing (bacteria and fungi) of 45 rock samples (sandstones) collected across 17 localities over 10 years of sampling activities in Antarctica with a 4-years microclimate monitoring based on *in situ* microclimatic sensing stations located in multiple locations, and information on rock physical and chemical properties. The results based on this extensive dataset allowed us to sort out, for the first time, those environmental conditions promoting life at the fringe of sustainability and those limiting the endolithic colonization in the Antarctic cold desert. Ecological networks were used to identify specific microbial assemblies (hereafter “Microbial assemblies” or “modules” or “clusters”) including groups of microbial species co-occurring in the same rocks. This approach allows us to simplify our analyses while working at the species level (phylotypes or zOperational Taxonomic Units (zOTUs) at 97% similarity), and further help us to identify the environmental preferences of different taxa, as taxa co-occurring in the same rocks should be sensible to similar environmental changes. We then used Random Forest (RF) analyses and non-parametric correlations to identify the major environmental drivers of Antarctic endolithic communities. Our new evidence fills an important knowledge gap that will definitely contribute to understanding the persistence of life at the edge on Earth, and suggests new strategies in the quest for extant or extinct life on other rocky planets such as Mars.

### Rock traits drive microbial diversity in Antarctica

Microbial endoliths represent the main life-form spreading in the Mars-like driest and coldest area known of the Antarctic Desert^3,8–9^. Besides, the capacity of rock substrates to support independent complex microbial communities is still far less understood. Using ecological network modeling, we identified ten independent microbial assemblies including co-occurring species of bacteria and fungi, thriving in distinct rock environments (Supplementary Figures S5, S6a, b; Supplementary Tables, sheets 3-4 to see the complete list of microbial phylotypes -species- in each assembly). This result indicates that a combination of different physic-chemical rock properties support the existence of multiple complex microbial assemblies at the edge of life. Thus, even under extreme conditions, such as those from Antarctica, environmental heterogeneity plays a fundamental role in supporting the coexistence of multiple microbial communities.

Using the Random Forest algorithm, we found that, compared with climate as mean annual temperature and mean annual precipitation and geographic features as elevation and distance from the sea,, rock traits are the most important factors controlling the proportion of bacteria and fungi within each microbial assembly, and further show that different combination of rock properties explain different microbial assemblies (Figure 1a). Rock diagenesis, quartz cement, sorting (i.e. distribution of grain size of sediments), and abundance of Iron (Fe) were the most important factors associated with microbial diversity, and with the proportion of different rock microbial assemblies (Figure 1a, 1b; Supplementary Figure S7). Among rock properties, rock porosity is widely recognized as the driving force of endolithic distribution and settlement^10–13^. Recent studies in Antarctica reported that the distribution of pores in sandstone was found to be more homogeneous compared to other rock typologies and allowed endolithic colonization to higher altitudes and greater distance from the sea^14,11^. Moreover, the physical structure, water-retention and capillarity capacity of the porous rocks helps to maintain moisture after the very rare precipitation events in the Dry Valleys, where water is inherently scarce^15^. Interestingly, rock porosity had an evident impact on bacterial richness, while it did not appear to affect fungi. This may due to the fact that bacteria, in particular *Cyanobacteria* can be affected by moisture content^16^, while fungi may outcompete other microbial groups under the most arid conditions, likely due to their high resistance to desiccation and great capacity to cope with drought stress and resurrect from dry conditions^17^.

**Figure 1.**
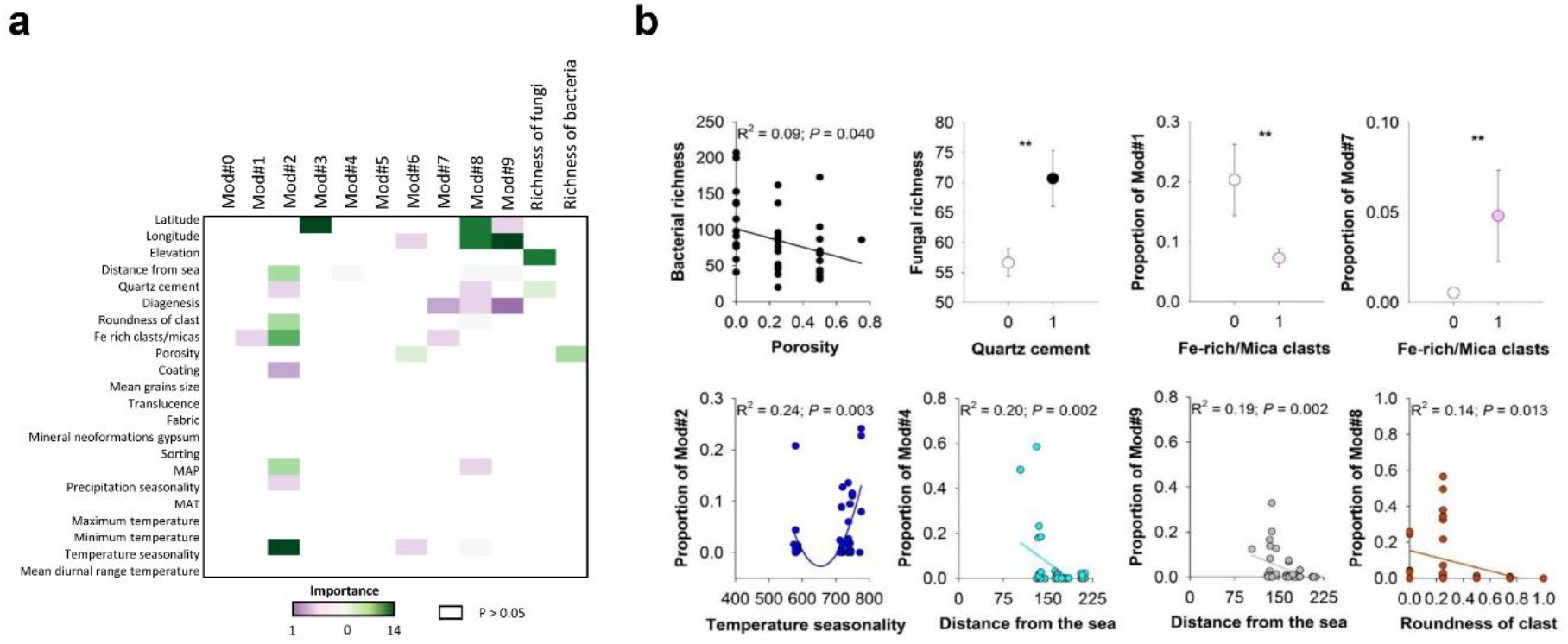
Most important factors driving the variation of Antarctic endolithic communities. **a)** Predictor importance (% increase in mean square error) from Random Forest modeling showing the most important drivers of the variation of microbial assemblies (Mod#) and fungal and bacterial richness. Importance measures with P < 0.05 (999 permutations). **b)** Selected examples showing the relationship between environmental factors and the proportion of modules and the richness of bacteria and fungi. Mod# = Microbial assembly.

Further chemical analyses in a subset of our rocks provided additional support to rock properties being essential drivers of contrasting microbial assemblies. For example, as shown in Figure 2, most of selected elements (Fe, S, K, P) correlates with Microbial assemblies#6 and #9, while Si and Ca did not show any significant correlation. Iron appeared as the most impacting elements, for which a correlation was evident in Microbial assemblies#1, #3, and #8, especially in the last two. Fe also strongly correlated with the bacterial richness, which, by the way, showed a high correlation with the other three elements too. In the area considered, sandstones belong to the Devonian to Triassic Beacon Supergroup, which mostly consists of quartzarenite, with Ferrar Dolerite intrusions of Jurassic age; the leaching and deflation of iron-bearing minerals from the overlying dolerite and its redistribution may contribute to increase Fe availability in the sampling sites^18^. Iron content is therefore high in such an environment and within local quartzarenite; iron-bearing minerals abundance clearly relates to biologically transformed iron. Fe-rich minerals (in the form of iron oxy-hydroxide microcrystals and Fe-bearing biogenic clays) are deposited around endolithic micro-inhabitants. Neoformed minerals are therefore accounted as biomarkers of previous endolithic activity^19^. Our results support the role of iron as a key-element in the geobiology of these communities. Beacon sandstone is rich in Si, almost consisting of quartz crystals; crystalline bulk silicon is rather inert, especially at low temperatures, but it confers translucence to rocks; the depth of PAR penetration is dependent on rock colour, mineralogy and structure^20^ and is fundamental to sustain phototrophic activity, which ultimately increases organic matter and feed the whole microbial community. Besides, in the present study no correlation with this element was found, maybe due to its indirect effect for the biological compartments considered (fungi and bacteria).

**Figure 2.**
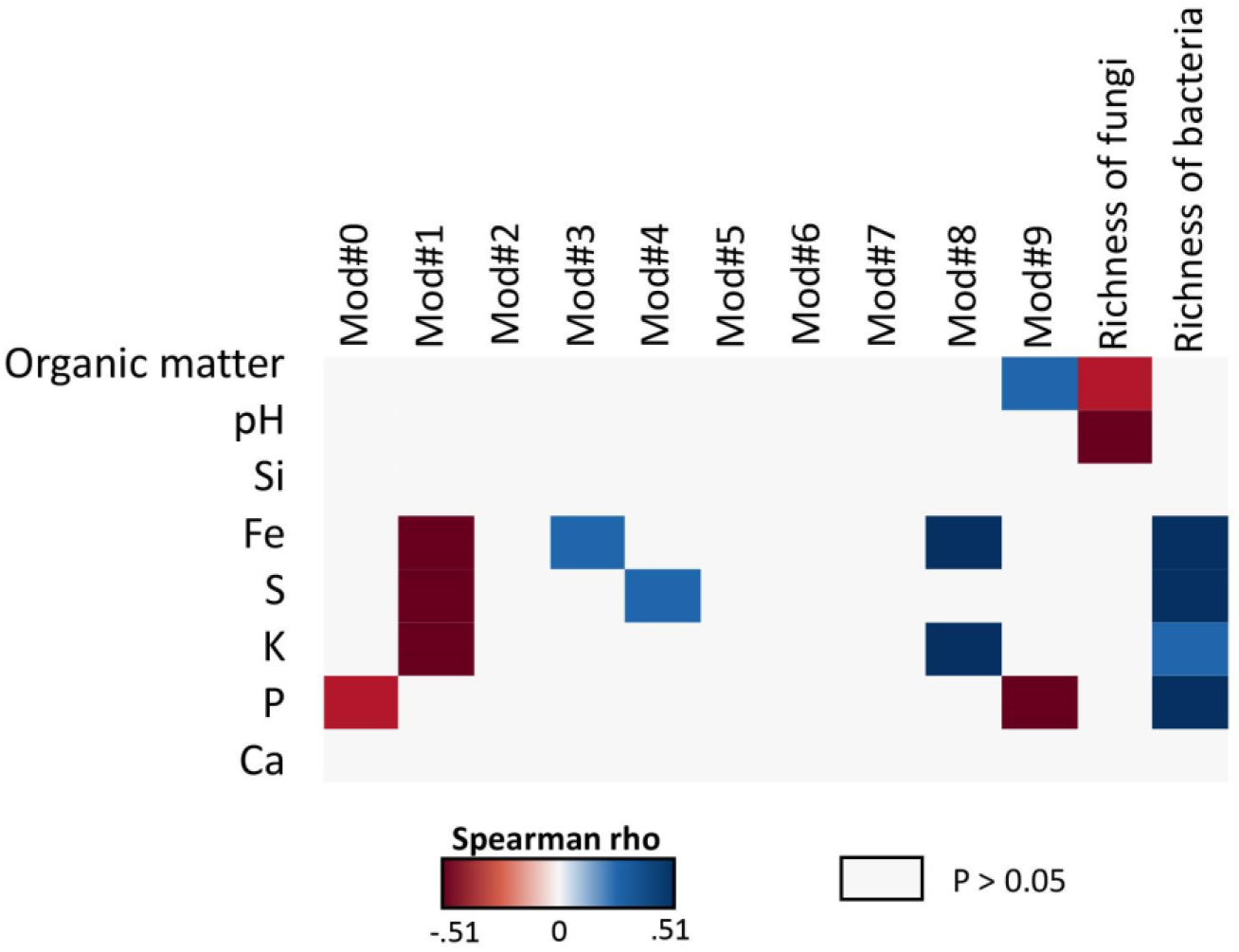
Correlations between biodiversity and physico-chemical data. Heatmap showing Spearman correlations between the relative abundance of modules and richness and rock properties. Mod# = Microbial assembly. White rectangles represent non-significant values (P > 0.05). Si = Silicon; Fe = Iron; S = Sulfur; K = Potassium; P = Phosphorus; Ca = Calcium.

The content of organic matter and pH have opposite effects on the richness of fungi and bacteria, respectively; fungi are included, as a whole, in a kingdom of purely heterotrophic organisms and are elective degraders in all environments. Moreover, they are very abundant in these communities that are, as a role, dominated by fungi, and lichens in particular. It is therefore not surprising that the organic matter correlated mainly with this biological compartment. For what concerns pH values, they exhibited a slight variation across the rocks analysed, ranging between 6.2 and 6.9. This range is optimal for most bacteria, which prefer neutral pH (6.5-7.5) and such a variation in their natural substrate may be ineffective. Differently, fungi are mostly a bit more acidophilic, preferring pH values ranging between 5 and 6; the pH variation outside their range preferences has therefore a more evident effect.

### Microclimate as critical factor in shaping taxa that are commonly associated with rock niche

Using low-resolution climatic data (Fig. 1), we found that climate can be important for certain microbial assemblies. To further investigate the role of climate in driving microbial assemblies, we conducted additional analyses using high-resolution microclimatic monitored data (see Methods). Microclimate (i.e. temperature of air, surface and inside rocks and PAR) played the role of major drivers of the relative abundance of three ecological clusters (Microbial assemblies#2, #5, and #9); however, the relative importance of these climatic factors was highly microbial assembly dependent (Figure 3). For instance, a weak, albeit significant, negative correlation between the relative abundance of Microbial assembly#2 was found, while the highest positive correlation was detected on Microbial assembly#9 (Figure 3a). This may be due to the fact that at genus level the bacterial and fungal phylotypes exhibited clear differences in their environmental preferences (and thus resulted in belonging to one or other microbial assembly), mainly ascribed to differences in climatic conditions (Supplementary Table, sheet 2).

**Figure 3.**
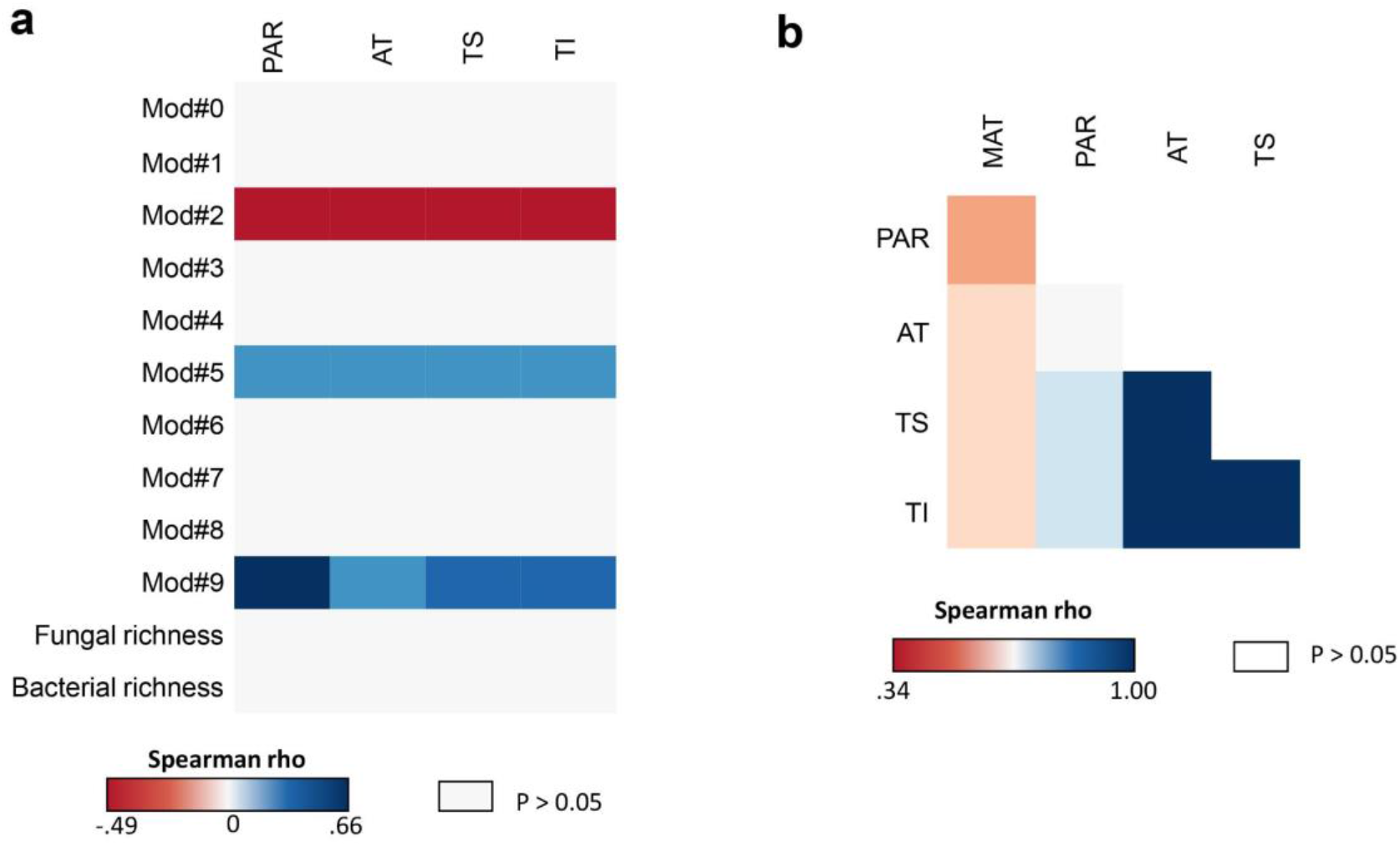
Correlations between biodiversity and microclimate. Heatmap showing Spearman correlations between the relative abundance of modules and richness and microclimatic monitored data from 2016 to 2019. Mod# = Microbial assembly. White rectangles represent non-significant values (P > 0.05). PAR = Photosynthetically active radiation. AT = Air temperature. TS = Temperature on rock surface. TI = Temperature inside rock.

Microbial assembly#9, in comparison with the other ecological clusters, contained higher diversity of taxa commonly and repeatedly associated with lithic niche, which have been isolated exclusively from Antarctic rocks. Specifically, microbial assembly#9 included fungal Operational Taxonomic Units (OTUs) belonging to the morpho-ecological group of black microcolonial fungi (or “Black Yeasts”) that encompass organisms specialized in poly-extreme-tolerance such *Cryomyces* spp. and *Coniosporium* spp.^21,22^. In particular, the endemic black fungus *Friedmanniomyces endolithicus* is among the most adapted and widespread having been retrieved in all Antarctic rock samples analyzed to date^8^. This microbial assembly contained also many basidiomycetes yeasts such as *Cryptococcus* spp., *Naganishia* spp., *Rhodotorula* spp.) with low temperature preferences, isolated also in other cold habitats such as Arctic permafrost^23^ and Alpine glaciers^24^. Microbial assembly#9 contained a large number of rock-bacterial taxa from the phylum Actinobacteria that represented the most abundant phylum in the bacterial genomes recently recovered from Antarctic endolithic metagenomes^7^. Remarkably, members of *Friedmanniella* spp. and *Conexibacter* spp., occurring in Microbial assembly#9, are typically recorded in highly oligotrophic environments such as cold-desert soils^25^.

Phylotypes from Bacteroidetes were also recorded in this Microbial assembly; in particular, the genus *Hymenobacter* that encompasses desert-bacteria with low temperature preferences such as *H. desertii*, *H. psychrophilus*, and *H. amundsei*, also isolated in rock samples from Antarctica^25,26^.

We also found that while PAR does not influence AT really, it has an effect on TS and TI: the intense solar radiation may heat the rock substratum and its thermal inertia may accumulate and maintain heating. MAT had a significant, albeit slight, effect on the other microclimatic parameters considered, while the most evident effect was the positive influence of temperature both of air and rock surface on temperature inside rocks (Figure 3b); this, together with the effect of PAR discussed above may significantly heating the endolithic niche and arguably activate microbial metabolism and growth.

In conclusion, our study provides unprecedented evidence of the enormous capacity of rocks and microclimatic combinations to support independent and complex microbial assemblies in challenging ecosystems from Antarctica. This first study clearly delineates the fundamental role of rock traits in driving their associated endolithic Antarctic communities. This is integral to understanding the fate of still biologically active hyper-arid drylands in a warming and drying world.

Ultimately, our work, identifying rock properties relevant for establishing the environmental conditions that promote microbial life on Earth, contribute to understanding patterns of terrestrial habitability and search for life during rover explorations on Mars surface.

## Methods

### Study area

The study area includes sandstones colonized by lichen-dominated cryptoendolithic communities from 21 locations in the Victoria Land (Continental Antarctica), which were collected during the XXVI (Dec. 2010–Jan. 2011) and XXXI (Dec. 2015–Jan. 2016) Italian Antarctic Expeditions (Supplementary Tables, sheet 1; Figure 4) in an altitudinal gradient from 834 to 3,100 m above sea level (a.s.l.), and ranging from −74.03 to −77-75 latitude and from 159.59 to 162.62 longitude. The sites included in the study have information available on the diversity of fungi and bacteria.

**Figure 4.**
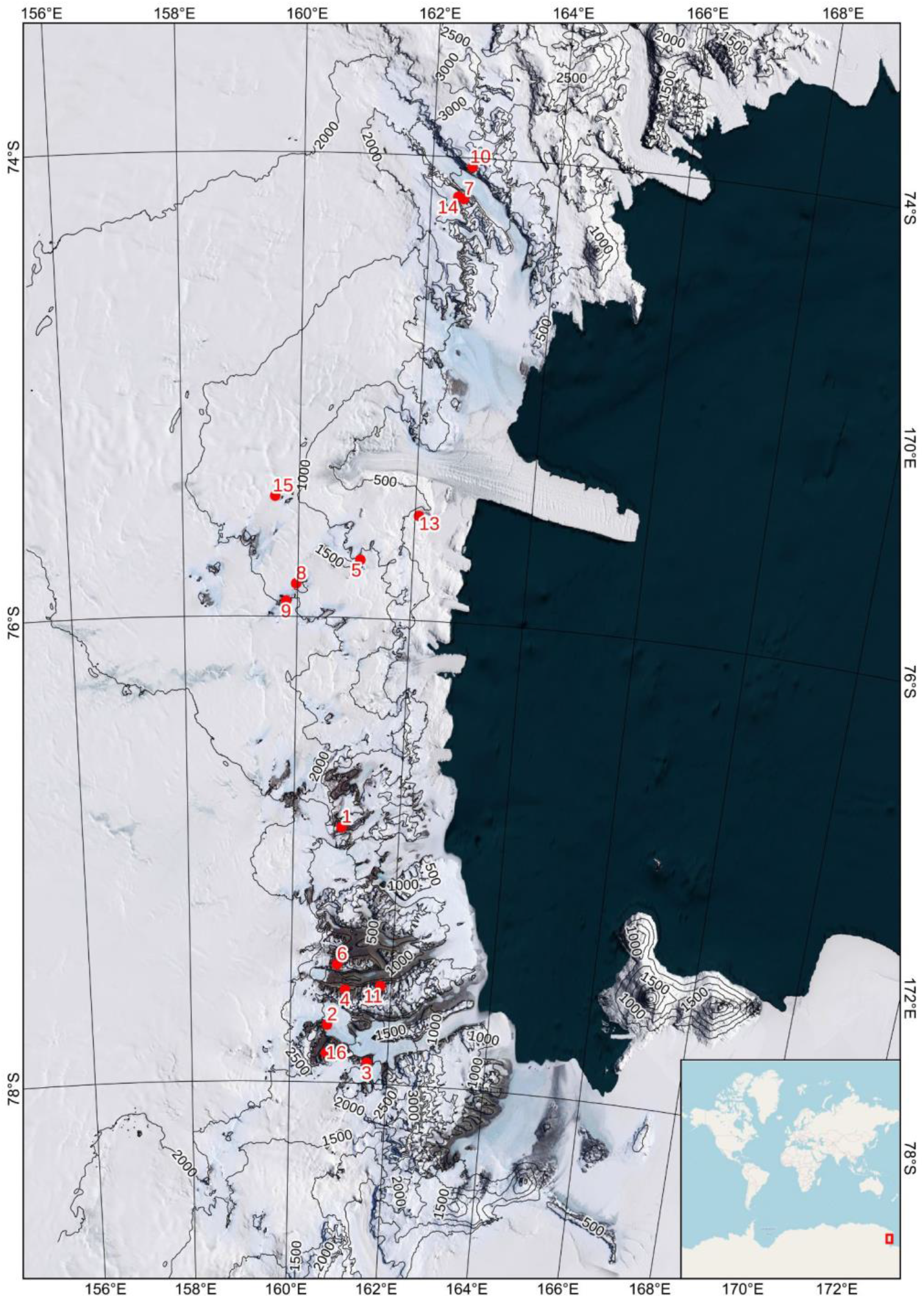
Map of the study area. Satellite imagery of the study area indicating elevations (contour lines) and the position of each sampling site; the inset reports the position of the study area in Antarctica. Key: 1) Battleship Promontory; 2) Finger Mt.; 3) Knobhead; 4) Linnaeus Terrace; 5) Mt. Bowen; 6) Mt. Elektra; 7) Mt. New Zealand; 8) Pudding Butte; 9) Richard Nunatak; 10) Shafer Peak; 11) Siegfried Peak; 12); The Mitten; 13) Thern Promontory; 14) Timber Peak; 15) Trio Nunatak; 16) University Valley.

Rocks (sandstones) were collected as described in^11^. Briefly, the presence of endolithic colonization (Supplementary Figure S1) was assessed by direct observation *in situ* and samples were aseptically excised using a geological hammer, collected in plastic sterile bags and transported, and stored at −20°C at the University of Tuscia (Italy), until DNA extraction and physic-chemical characterization.

### Molecular analyses

Illumina MiSeq was used for sequencing as described in^8^. Briefly, amplicons targeting the bacterial V4 region of 16S rRNA gene (F515/R806^27^), and fungal Internal Transcribed Spacer (ITS) 1 (ITS1F-ITS2^28^) were sequenced as reported in^21,29^. In all cases, global singletons and rare taxa (<5 reads) were eliminated as likely false positives due to sequencing errors, following^4,30^. Operational Taxonomic Units (OTUs) were clustered at 97% sequence similarity and OTU abundance tables were rarefied to ensure equal sampling effort across samples.

The Richness (i.e., number of OTUs) diversity index of each microbial group was calculated on these rarefied OTU tables (Figs. S2, S3).

All raw sequence data have been submitted to the GenBank databases under BioProject accession numbers PRJNA379160, PRJNA453198 and PRJNA379160 and to the European Nucleotide Archive (EMBL – EBI) under accession number PRJEB39480.

### Environmental and micro-climatic data

Mean annual temperature (MAT), mean annual precipitation (MAP), and precipitation seasonality (PSEA), were collected from the Worldclim database for all sites (https://www.worldclim.org; ∼1 km resolution^31^.

Microclimatic monitoring stations (HOBO USB Micro Station (ONSET Computer Corporation, Pocasset, MA) and microsensor (60 sec. resolution) were also installed across three notable sites to regularly record key parameters (water content, air temperature, surface temperature and temperature inside rocks, and Photosynthetic Active Radiation, PAR) for endolithic colonization were also installed in both northern (sun-exposed) and southern (shady) exposures across three notable sites: i) *Battleship Promontory*, BP, −76.90 160.91, 834 m a.s.l., McMurdo Dry Valleys, Southern Victoria Land; ii) *Pudding Butte*, PB, −75.85 159.97, 1573 m a.s.l., Northern Victoria Land; iii) *Mt. New Zealand*, MNZ, −74.18 162.51, 3100 m a.s.l., Northern Victoria Land).

Microclimatic data were acquired yearly from 2016 to 2019; for statistical analyses, we used microclimatic parameters averaged at the annual level, as explained below.

### Physicochemical analyses

Sandstones have been described for their general macroscopic properties and to assess the existence of external weathering rinds or coatings (Supplementary Figures S8-S11). As the reddish tone of most samples is related to the accumulation of Fe-bearing minerals, the color of each sample has been identified using the Munsell® code. Organic content and pH have been measured in the laboratory following as described in^32^. Briefly, total organic carbon (TOC) measurement required a titration using chromic acid to measure the oxidizable organic carbon^33^, while pH values have been measured with a pH meter (HI9321 – Hanna Instruments) after dispersing rocks in a large volume of pure water.

Sandstone rocks collected in the three monitored localities have been further investigated by microscopic analysis to assess weathering changes along a vertical transect. Rock samples were cut perpendicular to the topographic surface, polished, and thin section prepared according to the procedure described in^34^; this approach allowed to preserve the integrity of the external weathering surface and the presence of soluble minerals (salts and sulfates) ^32^. Micromorphological observation under plane-polarized light, cross-polarized light and oblique incident light of thin sections were conducted with an optical petrographic microscope Olympus BX41 equipped with a digital camera (Olympus E420); slides have been observed under oblique incident radiation (epifluorescence) of ultraviolet (UV) and blue (BLF) lights using a mercury burner (HBO lamp) 100 W^35^. Optical microscope permitted the description of the major properties of each sandstone outcrop (shape and roundness of grains, average size of grains, sorting, sedimentary structures, occurrence of siliceous cement, porosity, iron-rich or micaceous inclusions, diagenetic impregnation), the identification of mineral neo formations such as gypsum and oxalates^36^ and to assess the presence and lateral continuity of the external coating. Micromorphological studies of carbon coated thin sections also employed a Cambridge 360 scanning electron microscope (SEM) imaging both secondary and back-scattered electrons. Energy dispersive X-ray spectroscopy (EDS Link Isis 300) worked with an accelerating voltage of 20 kV, filament intensity 1.70 A, and probe intensity of 280 pA. Every analyzed element was previously standardized by using several single element standards provided by Micro-Analysis Consultants Ltd and elemental concentrations are reported as oxide weights normalized to 100%. In this study, we considered the concentrations of the most representative and abundant elements: Si, Fe, S, K, P and Ca. Such elements were selected because Si is the major constituent of the host rock; Fe is in charge of the color of the rock and Fe oxides are common in the coating of quartz grains; K and Ca are the most represented alkaline elements and are part of some of the mineralogical species identified as neoformations; P and S are elements whose origin is commonly related to biological processes and biomineralization^37,32^.

### Correlation network and statistical analysis

To identify clusters (modules or Microbial assemblies) of strongly associated rock taxa, a correlation network, i.e., co-occurrence network was performed. To produce a practicable correlation network, we kept bacterial and taxa exhibiting more than 80% of the relative abundance and then were then merged into a single abundance table. We then calculated all pairwise Spearman’s rank correlations (ρ) between all taxa. We focused exclusively on positive correlations to group all taxa with similar environmental preferences (i.e. OTUs co-occuring in the same locations). We considered a co-occurrence to be robust if the Spearman’s correlation coefficient was > 0.50 and *P* < 0.01^38^. The network was visualized with the interactive platform gephi^39^. The proportion of each module is calculated as the average of standardized abundances (0-1) of all OTUs within each microbial assembly (Supplementary Tables, sheet 4). By standardizing our data, we skip any effect of merging data from different groups (fungi and bacteria).

We then used the Random Forest (RF) model as described in^40^ to identify the major significant environmental predictors of rock diversity and the most important factors explaining the variation of modules and richness according to environmental variables and rock properties. The importance of each predictor variable is determined by evaluating the decrease in prediction accuracy, i.e., increase in the mean square error between observations and OOB predictions, when the data for that predictor is randomly permuted. RF was implemented using the ‘rfPermute’ package in R environment (http://cran.r-project.org/). Finally, we used Spearman correlations to further evaluate the relationship between environmental variables and rock properties and the richness of total fungi and bacteria and relative abundance of each microbial assembly.

## Acknowledgements

C.C. and L.S. acknowledge funding from the Italian National Program for Antarctic Research 478 (PNRA). C.C. is supported by a PNRA postdoctoral fellowship. M.D-B. is supported by a Ramón y 481 Cajal grant from the Spanish Ministry of Science and Innovation (RYC2018-025483-I).

## Author contributions

LS collected rock samples, CC, MD-B and LS developed the original idea of the analyses presented, interpreted results and wrote the first draft of the manuscript with the contribution of AZ, BT and PB, CC performed molecular and bioinformatic analyses, MD-B performed statistical, and AZ generated geological data. All authors have read and agreed to the published version of the manuscript.

## Competing interests

The authors declare no competing interests.

## Supplementary Tables legends

**Sheet 1:** Metadata of sandstone samples.

**Sheet 2:** Geomicrobiology analysis of sandstones collected in the three microclimatic monitored localities.

**Sheet 3:** Taxonomic assignment of bacterial and fungal taxa and ecological clusters (or Microbial assemblies) description.

**Sheet 4:** Relative abundance of each ecological cluster (or Microbial assembly) across the dataset.

## Notes

### Competing Interest Statement

The authors have declared no competing interest.

## References

1. Dragone, N. B. et al. Exploring the boundaries of microbial habitability in soil. J. Geophys. Res. Biogeosci. 126, (2021).

2. Torre, J. R. de la, de la Torre, J. R., Goebel, B. M., Imre Friedmann, E. & Pace, N. R. Microbial Diversity of Cryptoendolithic Communities from the McMurdo Dry Valleys, Antarctica. Applied and Environmental Microbiology vol. 69 3858–3867 (2003).

3. Archer, S. D. J. et al. Endolithic microbial diversity in sandstone and granite from the McMurdo Dry Valleys, Antarctica. Polar Biology vol. 40 997–1006 (2017).

4. Mezzasoma, A., Coleine, C., Sannino, C. & Selbmann, L. Endolithic Bacterial Diversity in Lichen-Dominated Communities Is Shaped by Sun Exposure in McMurdo Dry Valleys, Antarctica. Microbial Ecology (2021) doi:10.1007/s00248-021-01769-w.

5. Coleine, C. et al. Specific adaptations are selected in opposite sun exposed Antarctic cryptoendolithic communities as revealed by untargeted metabolomics. PLoS One 15, e0233805 (2020).

6. Onofri, S., Pacelli, C., Selbmann, L. & Zucconi, L. The Amazing Journey of Cryomyces antarcticus from Antarctica to Space. Extremophiles as Astrobiological Models 237–254 (2020) doi:10.1002/9781119593096.ch11.

7. Albanese, D. et al. Pre-Cambrian roots of novel Antarctic cryptoendolithic bacterial lineages. Microbiome vol. 9 (2021).

8. Coleine, C., Biagioli, F., Vera, J. P., Onofri, S. & Selbmann, L. Endolithic microbial composition in Helliwell Hills, a newly investigated Mars-like area in Antarctica. Environmental Microbiology (2021) doi:10.1111/1462-2920.15419.

9. Tyler, N. A. & Ziolkowski, L. A. Endolithic Microbial Carbon Cycling in East Antarctica. Astrobiology 21, 165–176 (2021).

10. Wierzchos, J. et al. Ignimbrite as a substrate for endolithic life in the hyper-arid Atacama Desert: Implications for the search for life on Mars. Icarus vol. 224 334–346 (2013).

11. Selbmann, L. et al. Effect of environmental parameters on biodiversity of the fungal component in lithic Antarctic communities. Extremophiles 21, 1069–1080 (2017).

12. Choe, Y.-H. et al. Comparing Rock-inhabiting Microbial Communities in Different Rock Types from a High Arctic Polar Desert. FEMS Microbiology Ecology (2018) doi:10.1093/femsec/fiy070.

13. Choe, Y.-H., Kim, M. & Lee, Y. K. Distinct Microbial Communities in Adjacent Rock and Soil Substrates on a High Arctic Polar Desert. Frontiers in Microbiology vol. 11 (2021).

14. Zucconi, L. et al. Mapping the lithic colonization at the boundaries of life in Northern Victoria Land, Antarctica. Polar Biology vol. 39 91–102 (2016).

15. Omelon, C. R. Endolithic Microbial Communities in Polar Desert Habitats. Geomicrobiology Journal vol. 25 404–414 (2008).

16. Starke, R. et al. Ecological and functional adaptations to water management in a semiarid agroecosystem: a soil metaproteomics approach. Scientific Reports vol. 7 (2017).

17. Gorbushina, A. A., Kotlova, E. R. & Sherstneva, O. A. Cellular responses of microcolonial rock fungi to long-term desiccation and subsequent rehydration. Studies in Mycology vol. 61 91–97 (2008).

18. Diaz, M. A. et al. Geochemistry of aeolian material from the McMurdo Dry Valleys, Antarctica: Insights into Southern Hemisphere dust sources. Earth and Planetary Science Letters vol. 547 116460 (2020).

19. Wierzchos, J., Ascaso, C., Sancho, L. G. & Green, A. Iron-Rich Diagenetic Minerals are Biomarkers of Microbial Activity in Antarctic Rocks. Geomicrobiology Journal vol. 20 15–24 (2003).

20. Walker, J. J. & Pace, N. R. Endolithic Microbial Ecosystems. Annual Review of Microbiology vol. 61 331–347 (2007).

21. Coleine, C. et al. Antarctic Cryptoendolithic Fungal Communities Are Highly Adapted and Dominated by Lecanoromycetes and Dothideomycetes. Front. Microbiol. 9, 1392 (2018).

22. Coleine, C. et al. Uncovered Microbial Diversity in Antarctic Cryptoendolithic Communities Sampling Three Representative Locations of the Victoria Land. Microorganisms vol. 8 942 (2020).

23. Faizutdinova, R. N., Suzina, N. E., Duda, V. I., Petrovskaya, L. E. & Gilichinsky, D. A. Chapter 8. Yeasts Isolated from Ancient Permafrost. Life in Ancient Ice 118–126 (2005) doi:10.1515/9781400880188-012.

24. Turchetti, B. et al. Influence of abiotic variables on culturable yeast diversity in two distinct Alpine glaciers. FEMS Microbiology Ecology vol. 86 327–340 (2013).

25. Pukall, R. et al. Complete genome sequence of Conexibacter woesei type strain (ID131577T). Standards in Genomic Sciences vol. 2 212–219 (2010).

26. Zhang, Q. et al. Hymenobacter xinjiangensis sp. nov., a radiation-resistant bacterium isolated from the desert of Xinjiang, China. International Journal of Systematic and Evolutionary Microbiology vol. 57 1752–1756 (2007).

27. Caporaso, J. G. et al. Ultra-high-throughput microbial community analysis on the Illumina HiSeq and MiSeq platforms. The ISME Journal vol. 6 1621–1624 (2012).

28. Smith, D. P. & Peay, K. G. Sequence depth, not PCR replication, improves ecological inference from next generation DNA sequencing. PLoS One 9, e90234 (2014).

29. Coleine, C. et al. Altitude and fungal diversity influence the structure of Antarctic cryptoendolithic Bacteria communities. Environ. Microbiol. Rep. 11, 718–726 (2019).

30. Lindahl, B. D. et al. Fungal community analysis by high-throughput sequencing of amplified markers – a user’s guide. New Phytologist vol. 199 288–299 (2013).

31. Zomer, R. J., Trabucco, A., Bossio, D. A. & Verchot, L. V. Climate change mitigation: A spatial analysis of global land suitability for clean development mechanism afforestation and reforestation. Agriculture, Ecosystems & Environment vol. 126 67–80 (2008).

32. Zerboni, A. Holocene rock varnish on the Messak plateau (Libyan Sahara): Chronology of weathering processes. Geomorphology vol. 102 640–651 (2008).

33. Walkley, A. & Armstrong Black, I. An examination of the degtjareff method for determining soil organic matter, and a proposed modification of the chromic acid titration method. Soil Science vol. 37 29–38 (1934).

34. Fitzpatrick, E. A. Thin Section Preparation of Soils and Sediments. By C. P. Murphy. Berkhamsted, UK: AB Academic Publishers (1986), pp. 149, £18.95 (paperback). Experimental Agriculture vol. 23 472–472 (1987).

35. Stoops, G. Fluorescence Microscopy. Archaeological Soil and Sediment Micromorphology 393–397 (2017) doi:10.1002/9781118941065.ch36.

36. Cremaschi, M., Trombino, L. & Zerboni, A. Palaeosoils and Relict Soils. Interpretation of Micromorphological Features of Soils and Regoliths 863–894 (2018) doi:10.1016/b978-0-444-63522-8.00029-2.

37. Reneau, S. L., Raymond, R. & Harrington, C. D. Elemental relationships in rock varnish stratigraphic layers, Cima volcanic field, California; implications for varnish development and the interpretation of varnish chemistry. American Journal of Science vol. 292 684–723 (1992).

38. Barberán, A. The microbial contribution to macroecology. Frontiers in Microbiology vol. 5 (2014).

39. Bastian, M., Heymann, S. & Jacomy, M. Gephi: An Open Source Software for Exploring and Manipulating Networks. ICWSM 3, 361–362 (2009).

40. Delgado-Baquerizo, M. et al. Ecological drivers of soil microbial diversity and soil biological networks in the Southern Hemisphere. Ecology 99, 583–596 (2018).

