## Supporting Data for "Rock traits drive complex microbial communities at the edge of life"

**Supplementary Materials for the article entitled: Rock traits drive complex microbial communities at the edge of life**

This file includes:

Supplementary Figures S1-S11

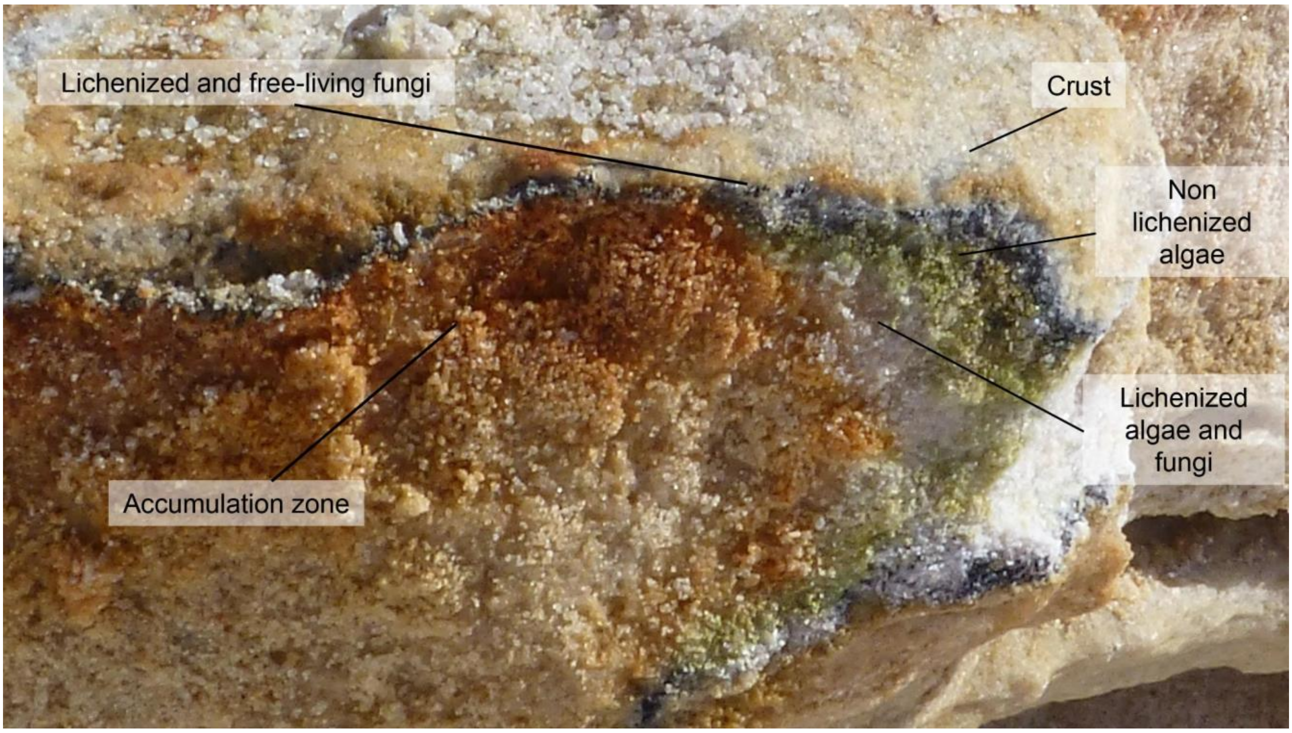


**Figure S1.** Cryptoendolithic lichen-dominated community colonizing a sandstone sample at Linnaeus Terrace, McMurdo Dry Valleys, Southern Victoria Land.

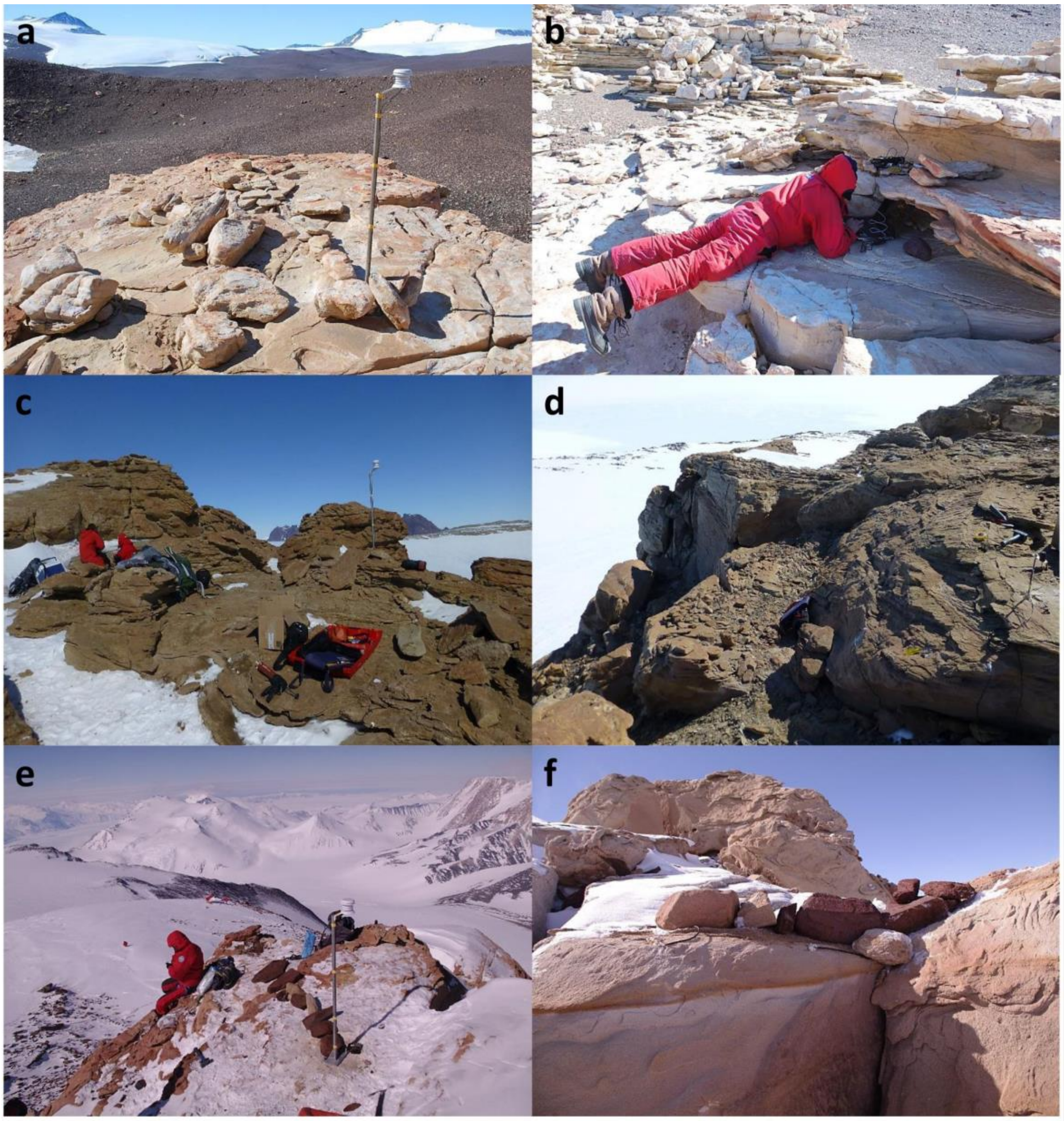


**Figure S2. Microclimate monitoring stations.** Microclimatic monitored stations installed in both north sun exposed and south shady rock surface: **a-b)** Battleship Promontory (834 m a.s.l., McMurdo Dry Valleys, Southern Victoria Land) north and south, respectively; **c-d)** Pudding Butte (1,573 m a.s.l., Northern Victoria Land) north and south, respectively; **e-f)** Mt. New Zealand (3,100 m a.s.l., Northern Victoria Land) north and south, respectively

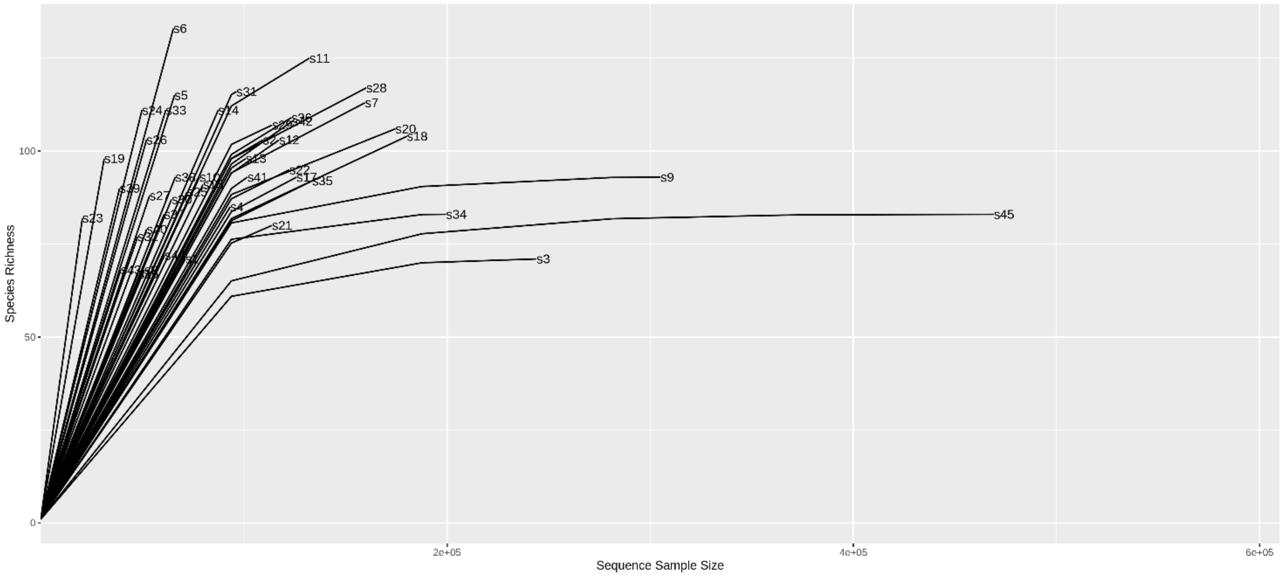


**Figure S3.** The frequency of observed fungal Operational Taxonomic Units (OTUs) for each sample was used to calculate fungal rarefaction curves.

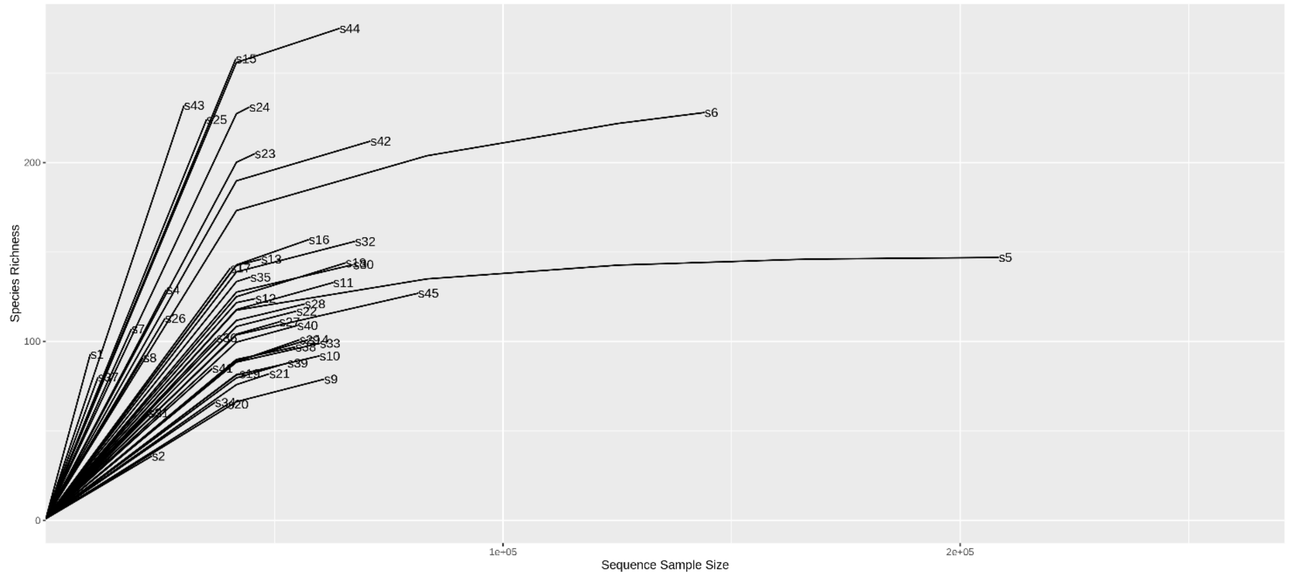


**Figure S4.** The frequency of observed bacterial Operational Taxonomic Units (OTUs) for each sample was used to calculate bacterial rarefaction curves.

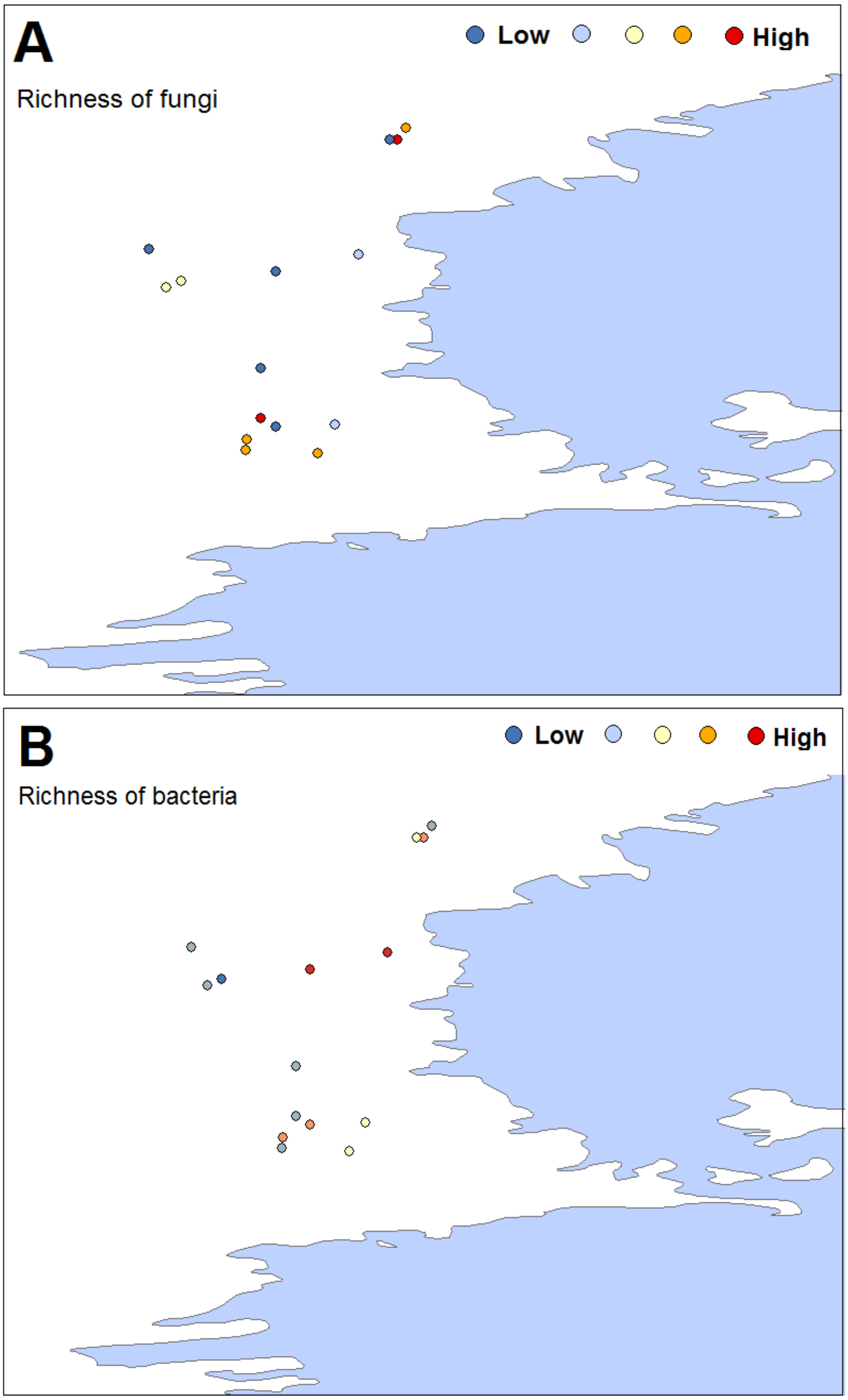


**Figure S5.** Richness of fungi **(a)** and bacteria **(b)** across the study area. Circles represent the sampled sites.

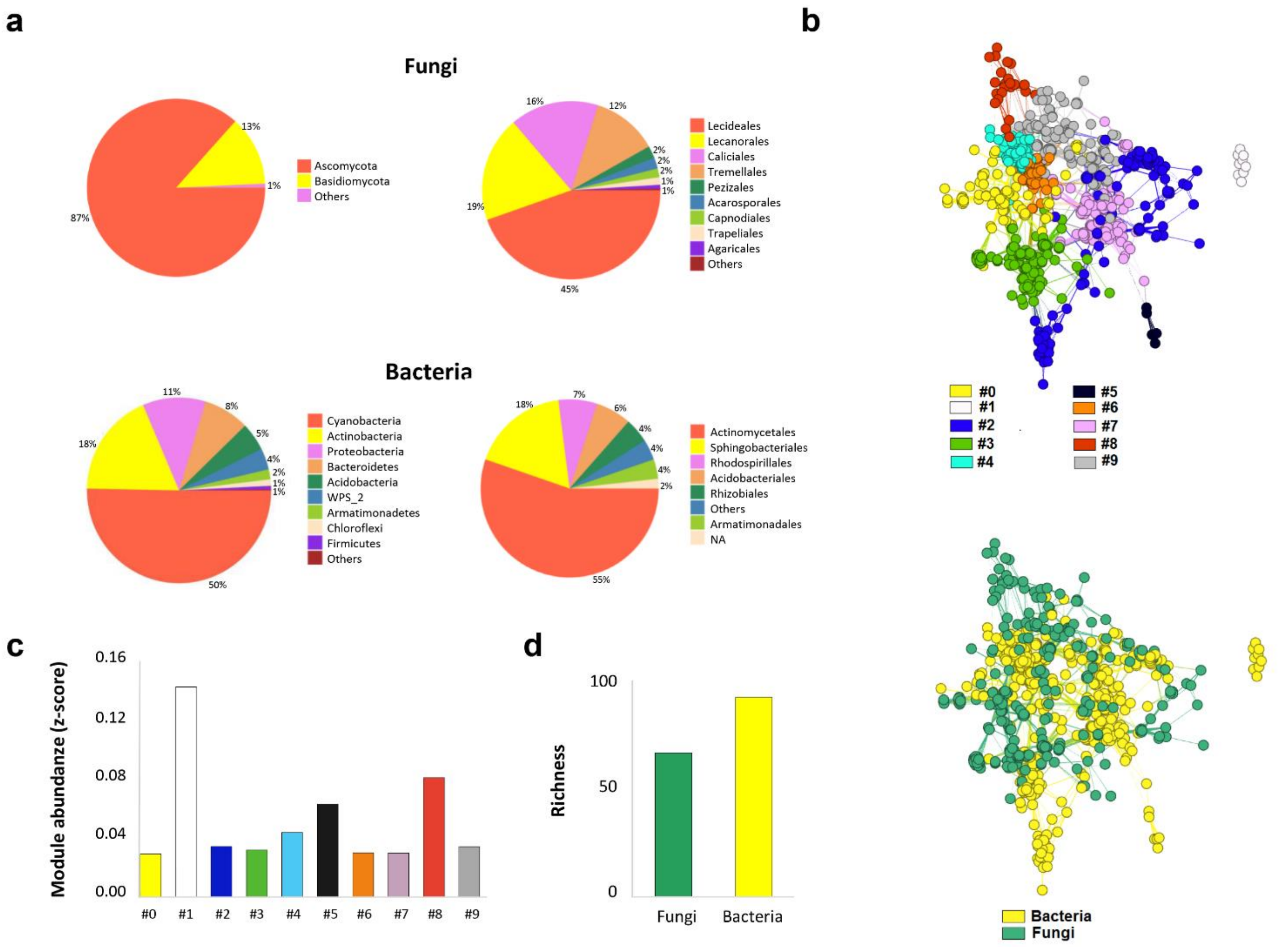


**Figure S6. a)** Relative abundance of main groups of fungi and bacteria in the surveyed sites. **b)** Rock correlation networks representing a network diagram with nodes (taxa of fungi and bacteria). Different colors represent the different identified modules (first network) and those belonging to fungi or bacteria (second network). **c)** Histogram showing the mean values of each module. **d)** Histogram showing the mean values of total fungi and bacteria.

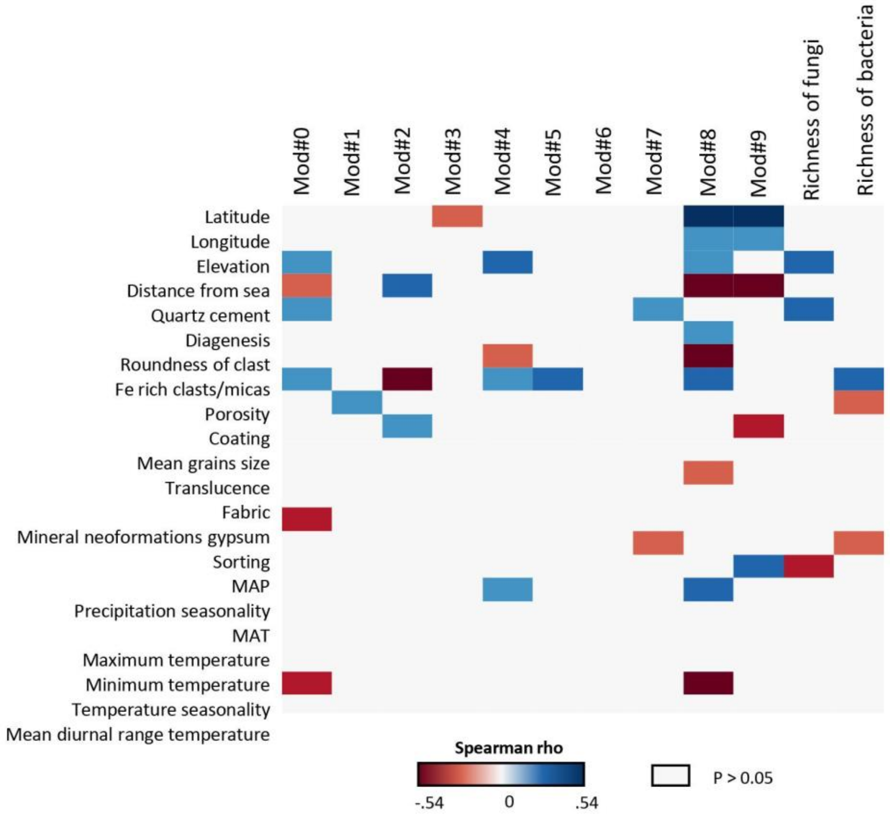


**Figure S7.** Heatmap showing Spearman correlation between environmental variables and rock properties and relative abundance of modules and richness ($P < 0.05$).

White rectangles represent non significant values.

MAP = abundance and mean annual precipitation.

MAT = mean annual temperature.

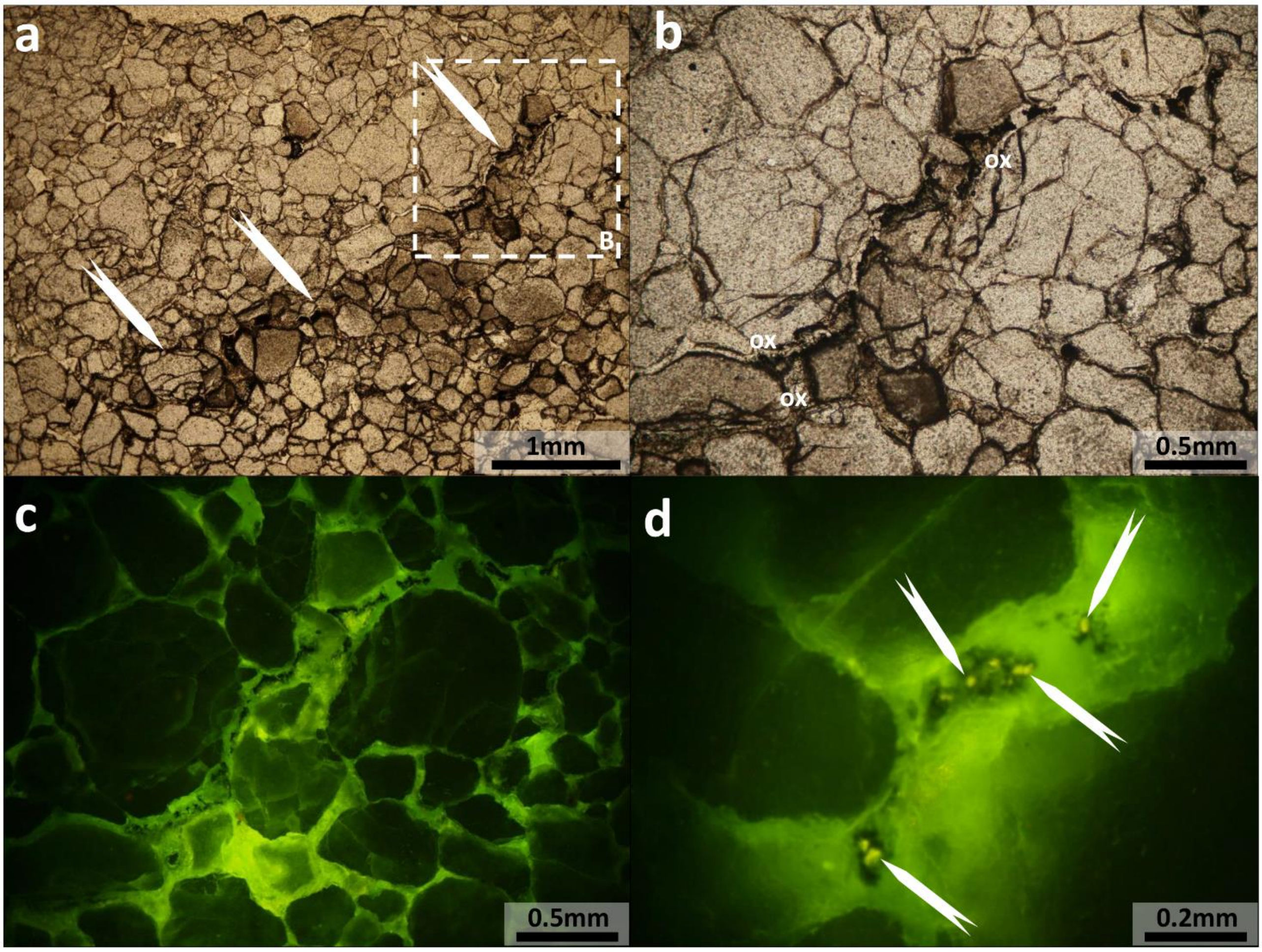


**Figure S8. Photomicrographs of sandstone from sampling site Battleship Promontory. a)** General view in plain polarized light of near-surface sandstone; note the planar discontinuity infilled by dark material (indicated by the arrows). **b)** A detail of the planar discontinuity of (a), infilled by dark organic particles and bright micro-crystals of oxalates (ox). **c)** The same of (b) under incident radiation of blue light suggesting the occurrence of biogenic minerals and phosphatic material. **d)** A detail of the organic matter-rich constituents including phosphatic crystals (indicated by the arrows).

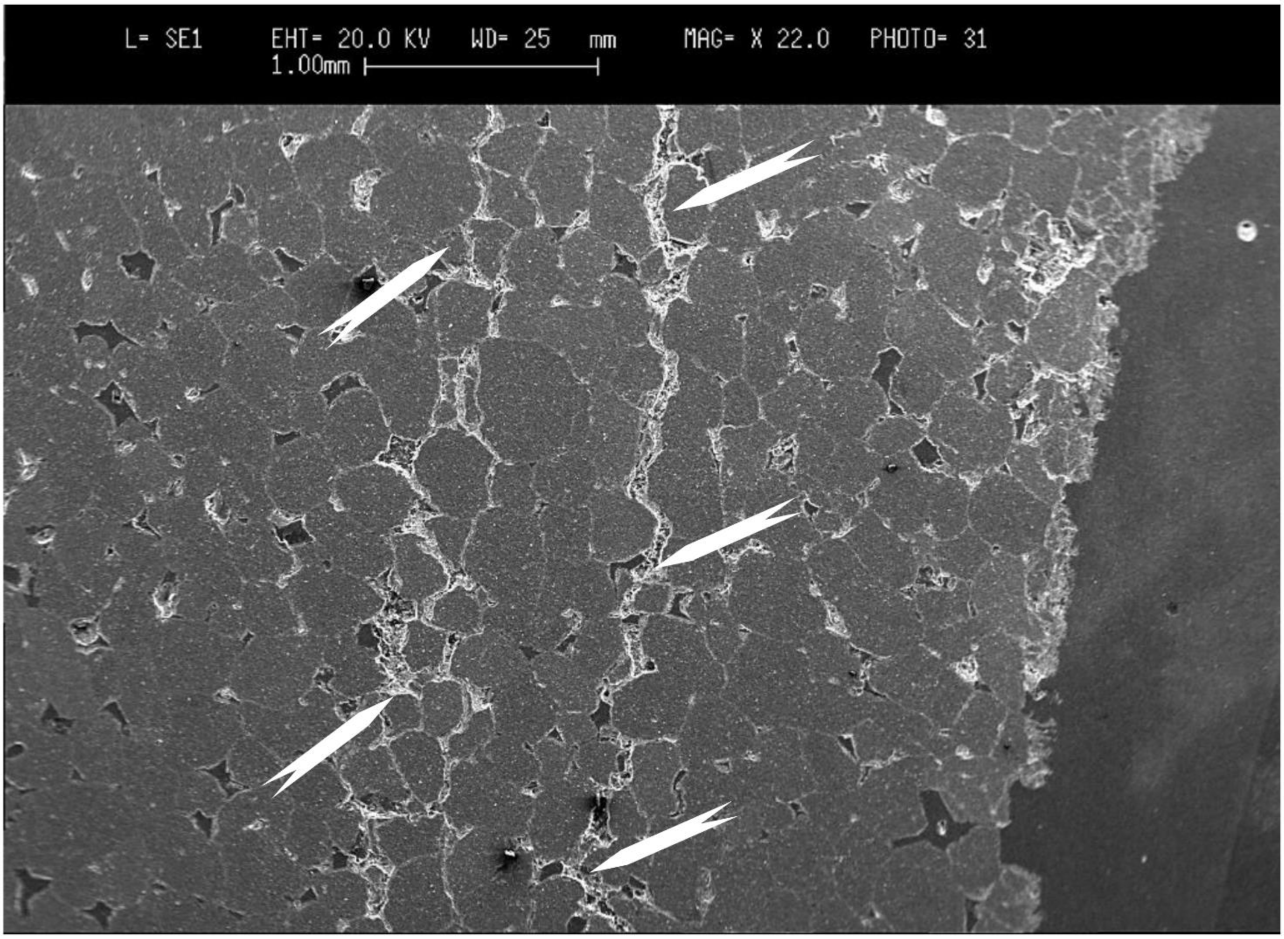


**Figure S9.** Scanning Electron Microscope imaging of near-surface sandstone illustrating the occurrence of planar discontinuities (indicated by the arrows) infilled by accumulation of oxalates.

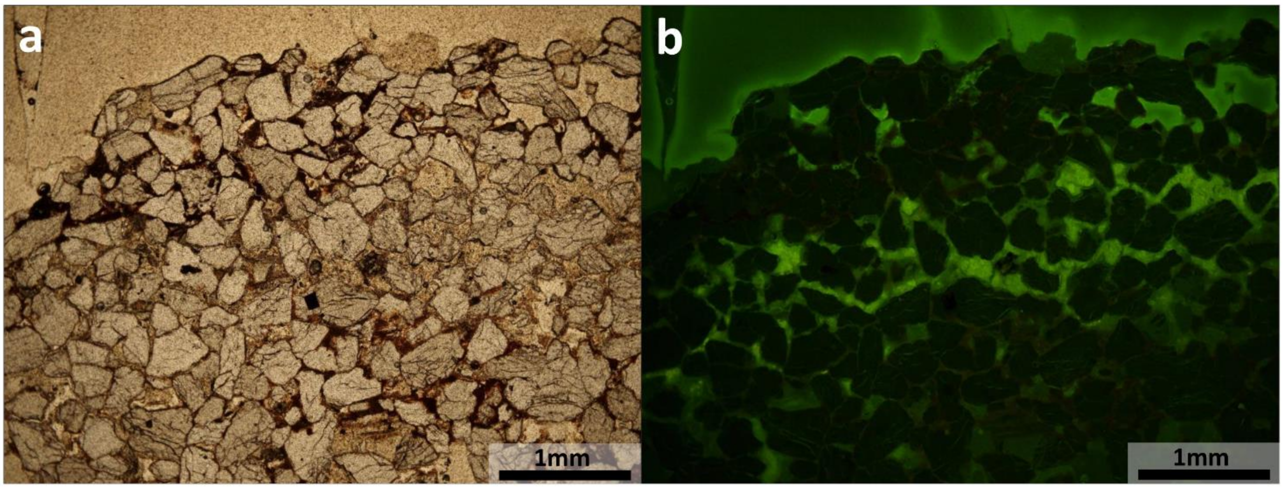


**Figure S10. Photomicrographs of sandstone from sampling site Pudding Butte. a)** The picture illustrates the general aspect of local sandstone that display abundant secondary infilling in the near-surface part. **b)** The same under incident radiation of blue light indicating the occurrence of biogenic minerals only in the outer part of the sample.

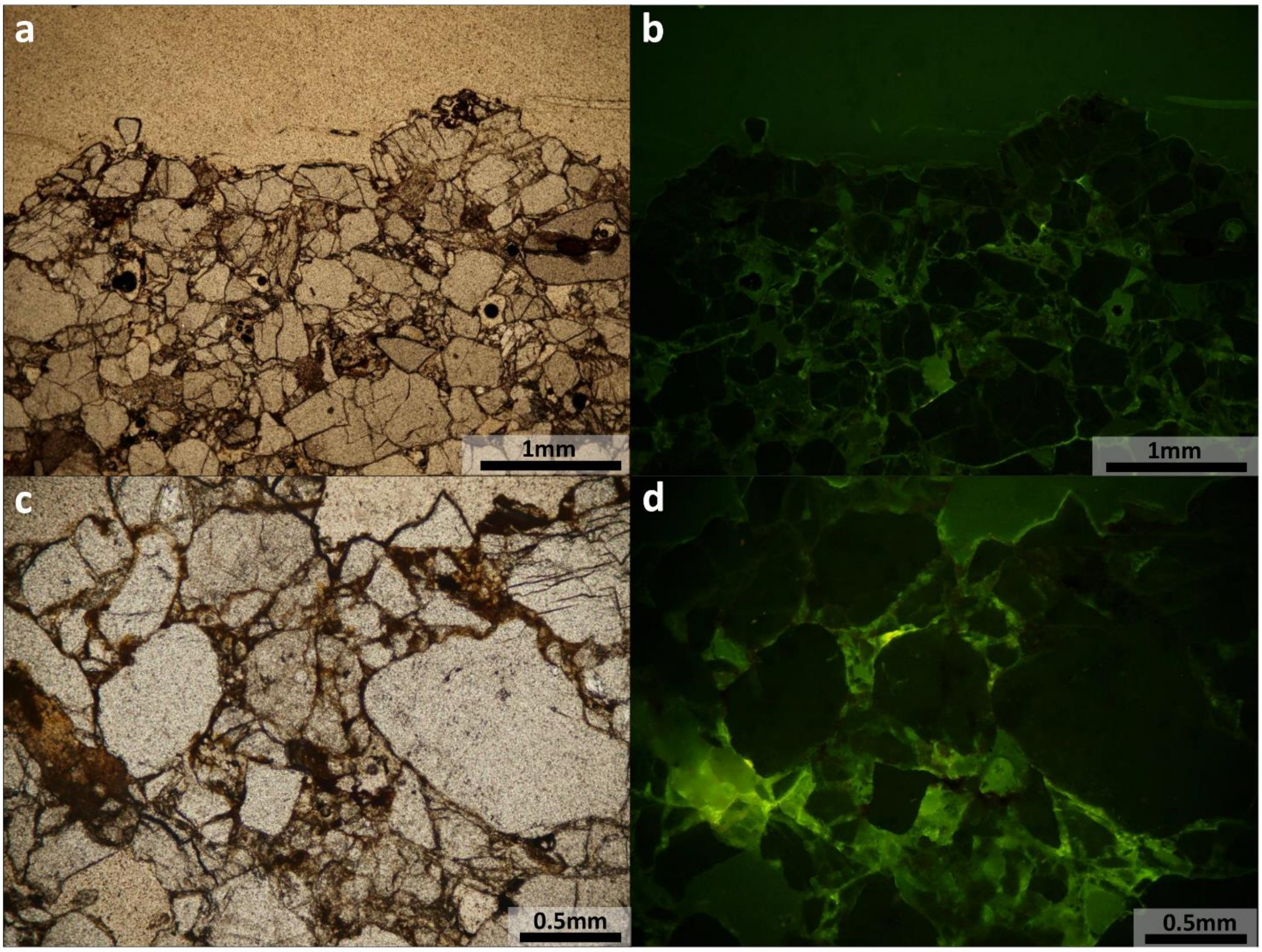


**Figure S11. Photomicrographs of sandstone from sampling site Mt. New Zealand. a)** General view of the sandstone including Fe-rich minerals and clay under plain polarised light. **b)** The same of (a) under epifluorescence of blue light. **c)** A detail of (a). **d)** The same under incident radiation of blue light suggesting the occurrence of biogenic minerals.
